# TITINdb2 – Expanding Annotation and Structural Information for Protein Variants in the Giant Sarcomeric Protein Titin

**DOI:** 10.1101/2024.05.08.593166

**Authors:** T. Weston, J. C-F. Ng, O. Gracia Carmona, M. Gautel, F. Fraternali

## Abstract

**Summary:** We present TITINdb2, an update to the TITINdb database previously constructed to facilitate the identification of pathogenic missense variants in the giant protein titin, which are associated with a variety of skeletal and cardiac myopathies. The database and web portal have been substantially revised and include the following new features: (i) an increase in computational annotation from 4 to 20 variant impact predictors, available through a new custom data table dialogue; (ii) thorough structural coverage of single domains with AlphaFold2 predicted models; (iii) newly predicted domain-domain interface annotations; (iv) an expanded *in silico* saturation mutagenesis incorporating 4 variant impact predictors; (v) a comprehensive overhaul of available data, including population data sources and variants reported pathogenic in the literature; (vi) A curated mapping of existing protein, transcript and chromosomal sequence positions and a new variant conversion tool to translate variants in one format to any other format.

**Availability and Implementation:** Database accessible via titindb.kcl.ac.uk/TITINdb/

**Contact:** Franca Fraternali

**Supplementary Information:** Available

## Introduction

The wealth of information provided by next-generation sequencing technologies in mapping the landscapes of genetic variation across populations have provided valuable insight into the tolerance of their protein products to specific mutations; however, elucidation of the precise impact of a given variant remains both challenging and costly. This issue is especially notable in proteins such as titin, where the sheer size of the protein – 35991 amino acid residues across 363 exons (Maruyama 1976; Labeit *et al*. 1990, 1992) – precludes experimental characterisation of every possible variant and renders the probability of finding rare, possibly pathogenic variants even in healthy individuals a near certainty. Thus, the characterisation of damaging variants in titin, most notably skeletal and cardiac myopathies, is both time- and labour-intensive. Computational methods represent an efficient solution for prioritising these investigations.

To address these issues, the web server TITINdb was established (Laddach, Gautel and Fraternali 2017), which allows the user to visualise any residue in the 302 titin domains (169 immunoglobulin, 132 fibronectin type-III, 1 protein kinase) on an atomic-level experimental structure or model of that domain, and to extract information on its contacts, solvent accessibility, and the frequency of observed variation at that position. Furthermore, this functionality was integrated with both population variant data and reported pathogenic variants in the literature, with the goal of facilitating the prioritisation of candidates for subsequent experimental characterisation. Since its release, new technologies and approaches have been developed, such as large language models for protein structural modelling (Jumper *et al*. 2021) and variant impact prediction (Cheng *et al*. 2023a), alongside further expansion of the existing genetic data sets of both population and clinically relevant single amino acid variants (SAVs). We have therefore updated TITINdb to take advantage of this wealth of information, and to promote the continued improvement of our understanding of the consequences of titin SAVs.

## TITINdb2 Features and Data

### Data Sources

The original release of TITINdb included known SAVs in titin from three sources: variants reported as pathogenic in the literature, population variants from the Genome Aggregation Database (gnomAD) (Lek *et al*. 2016; Karczewski *et al*. 2020) and population variants from the 1000 Genomes Project (1KGP) (The 1000 Genomes Project Consortium 2015). TITINdb2 expands the data collection to include three gnomAD data sets (gnomAD-2.1.1, gnomAD-3.1.2, and gnomAD-4.0.0) as separate categories. Thus, in total TITINdb2 contains 48,288 distinct, annotated missense variants, of which 76 variants are reported pathogenic in the literature. To obtain these reported variants, we collected published evidence and case studies in which titin missense variants were identified (Weston *et al*. 2024).

### AlphaFold2 and AlphaMissense – Structural Coverage and Variant Annotation

The structural coverage of titin domains has been supplemented by the release of AlphaFold2 predictions for the human proteome (Jumper *et al*. 2021); however, titin as a complete protein is too large to be represented in a single model, and thus these high-quality models of titin domains are not readily available. Thus, we have extracted the full predictions for titin from the complete human proteome predictions available at AlphaFoldDB (AlphaFold Protein Structure Database); here, the structure of the canonical sequence of titin is predicted in 166 overlapping “contigs” of 1400 residues (see Supplementary Material). The structures of individual titin domains were extracted from each contig according to the domain boundaries established in the previous release of TITINdb (Laddach, Gautel and Fraternali 2017). For each individual domain, we obtained an ensemble of 6-7 predicted models from these contigs. We additionally include tandem Ig domains, where two Ig or Fn3 domains are connected by a short (1-8 residue) linker, as separate structures.

TITINdb2 offers rapid access to the curated AlphaFold2 domain structures alongside quality assessment metrics (Rama Z-score, MolProbity (Williams *et al*. 2018) score) and mapping of SAVs to these structures. Variant impact scores predicted by the recently published AlphaMissense (Cheng *et al*. 2023a), a model based on AlphaFold2, have been extracted from proteome-wide SAV impact predictions (Cheng *et al*. 2023b) and given as annotations in TITINdb2. Additionally, we have added a functionality to filter variants by aligned position within homologous titin domains, allowing for hotspots specific to domain folds to be better recognised; for an example of this approach, see Case Study 2 in the Supplementary Material.

### Domain-Domain Interface Annotation

A newly added feature in TITINdb2 is the annotation of domain-domain interface residues. This is of relevance to the study of titin as the protein *in vivo* is composed of a chain of many closely linked domains, such that residues located at the interfaces between two titin domains may be buried by other residues in the linker or adjacent domain. Thus, annotation of solvent accessibility based on analysis of single-domain models may not be appropriate for residues at these interface positions.

Thanks to the provision of AlphaFold2 structural models, we now have models of all tandem two-domain constructs in the whole titin protein, and thus we have introduced annotations of these interface residues to TITINdb2. In brief, a residue is labelled as “interface-buried” based on the difference between the solvent-accessible surface area (SASA; calculated by POPS (Cavallo, Kleinjung and Fraternali 2003)) for a residue in the domain alone and that same residue in the multi-domain tandem. The predicted interface-buried residues are largely consistent between different AlphaFold2 predicted models, and our analysis indicates that 7.4% of domain residues in titin are interface-buried by this definition (for further details, see Supplementary Material).

### Annotation with new Variant Impact Predictors

In addition to the annotation of SAVs provided in the original TITINdb with Condel (González-Pérez and López-Bigas 2011), mCSM (Pires, Ascher and Blundell 2014a), DUET (Pires, Ascher and Blundell 2014b), and PolyPhen2 (Adzhubei, Jordan and Sunyaev 2013), we have updated the database with a broader range of variant impact predictors. Notably, we have computed pathogenicity prediction scores for all possible SAVs with Rhapsody (Ponzoni *et al*. 2020), for all domains in the canonical N2BA isoform (i.e. excluding domains Ig25, Ig26), using as input predictions retrieved from PolyPhen2. We have also extracted REVEL (Ioannidis *et al*. 2016) scores from the online genome segment files (REVEL genome segment files) for all possible nsSNVs in titin. Thus, we have computational saturation mutagenesis predictions calculated from seven tools: AlphaMissense, Condel, DUET, PolyPhen2, mCSM, REVEL, and Rhapsody, as well as pre-computed pathogenicity prediction scores from 13 tools sourced from dbNSFPv4 (Liu *et al*. 2020). All the available predictors can be selected or de-selected in the visible table using the new Custom Data Table functionality. A full list and information on the predictors included can be found in the Supplementary Material.

### TITINdb2 Variant Converter

A commonly reported issue for scientists working with titin is that of the difficulty in converting between protein, transcript, and genomic coordinates, for which a tool is not readily available; as titin is composed of multiple isoforms and the gene is coded on the reverse strand, this is especially prominent. In TITINdb2, we have conducted a comprehensive mapping between these coordinates in the seven major isoforms of titin (IC, N2AB, N2B, N2A, novex-1, novex-2, novex-3) to the corresponding transcript and chromosomal sequences. These data have been added to every “Position” page in TITINdb2, alongside wild-type codons for both transcript and genomic sequences. In addition, we have designed and built a “Variant Converter” application accessible through TITINdb2, which can convert input variants in bulk from any isoform and format to any other, greatly simplifying this process for the end user. Further information may be found in the Supplementary Material.

## Applications

We here briefly describe an example case study showcasing the new features implemented in TITINdb2. Further details, including figures and additional case studies, may be found in the Supplementary Material.

### Comparing Variant Impact across Positions and Populations

While annotations of individual variants give an idea of the impacts and characteristics of that specific variant, they must be contextualised by comparison to other variants in the same and other domains. While experimental saturation mutagenesis methods to view the full mutational landscape of a protein are useful, these are limited by the constraints of time and expense (Fowler and Fields 2014), and thus are hard to apply to giant proteins such as titin. Alternatively, *in silico* saturation mutagenesis offers a way to assess the predicted impact of variants, using computational variant assessment tools; however, there are many different possible methods, each with their own advantages and disadvantages. In TITINdb2, seven predictive tools have been used to annotate either the whole or the majority of the possible protein substitutions in titin; included in this number is AlphaMissense (Cheng *et al*. 2023a), a new and potentially very powerful pathogenicity prediction tool for which pre-computed predictions for many proteins, including titin, have been made available to the scientific community.

To investigate how these annotations compared when applied to known SAVs, we annotated all reported domain-located titin SAVs in population and disease-associated variant databases with the predicted pathogenicity score from AlphaMissense, as well as two generally well-regarded conventional predictors: Rhapsody, a predictor incorporating a wide array of sequence, structural, and dynamical features; and REVEL, a meta-predictor reporting a consensus score from a number of constituent sequence-based predictors. For Rhapsody, we used the AlphaFold2 predicted structures available in TITINdb2 as inputs for predictions, and for REVEL, we used TITINdb2’s variant converter tool to extract predictions from the available genome segment files and convert to protein coordinates. All three of these predictors report a score from 0 to 1, where 0 indicates a high-likelihood neutral variant, and 1 a high-likelihood pathogenic variant.

All three predictors were able to identify most of the 33 experimentally validated pathogenic variants as damaging (Figure 1d); however, AlphaMissense predicted most of these variants with high certainty, whereas both REVEL and Rhapsody had more uncertainty in their predictions. Similarly, the distribution of predictions over variants in the population genetic databases gnomAD-4.0.0, gnomAD-3.1.2 and gnomAD-2.1.1 differed between the three predictors: AlphaMissense returned a bimodal distribution of predictions, where most variants were predicted with high certainty to be either pathogenic or neutral; Rhapsody returned a more spread distribution, with the fewest variants in high-certainty bins; REVEL returns a skewed distribution where most population variants are considered neutral (Figure 1c).

**Figure 1.**
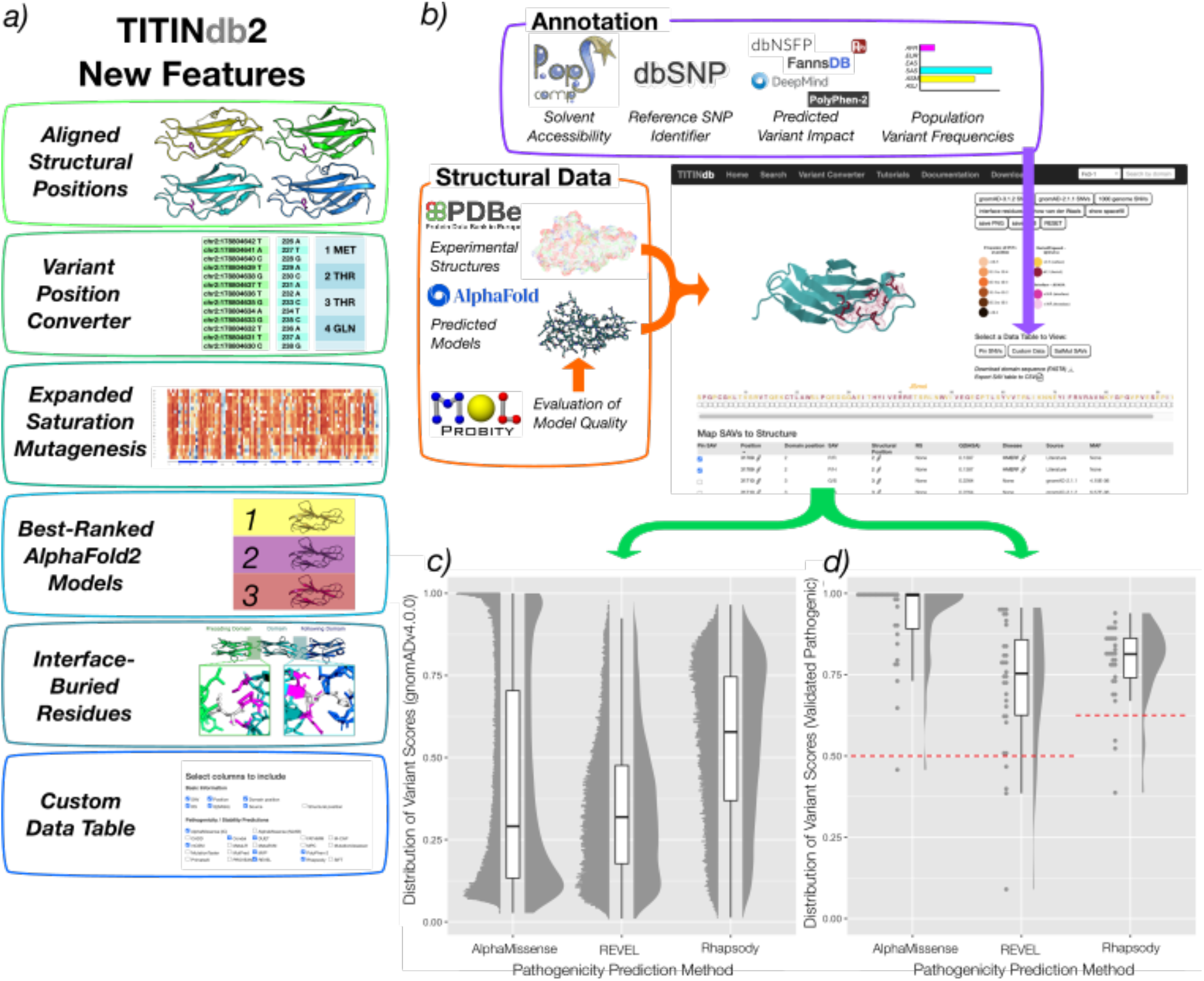
a) Highlighted new features available in TITINdb2 not present in the previous version. Access to Alphafold2 predicted domain models has allowed for comparative metrics to other models, aligned structural positions, and interface-buried residues. Increased variant impact prediction annotations and saturation mutagenesis may be stratified according to the user’s needs by the Custom Data Table option. b) Screen capture of TITINdb2, showing variants on titin Fn3-119 with AlphaFold2 predicted structure. Annotated to show different sources of data collated in database and data layout in page. Variants reported in c) the recently released gnomAD-4.0.0 population variant database and d) disease-associated variants with experimental validation are annotated with three commonly used variant impact predictors, AlphaMissense, REVEL and Rhapsody, and plotted as stacked bubble, box, and violin plots to visualise the distribution of predictions across these data sets.

These findings have implications for the use of each of these tools in practical applications with respect to titin. AlphaMissense predictions are the most capable of successfully identifying pathogenic variants and do so with high certainty; however, they also indicate a larger-than-expected quantity of population genetic variants as similarly high likelihood of pathogenicity, suggesting issues with false positives. REVEL predictions are the weakest at identifying these pathogenic variants but identify almost all population variants as neutral in keeping with our assumptions. Rhapsody predictions give more information than either REVEL or AlphaMissense through a broader spectrum of predicted damage, but also misclassify more pathogenic variants than AlphaMissense and more population variants than REVEL. As such, each of these tools has strengths that lend them to better performance in different research contexts; AlphaMissense for identifying potentially pathogenic variants of interest, REVEL for identifying background variation, and Rhapsody for delineating between greater and lesser damaging variants.

## Acknowledgements

The authors would like to thank those who have helped bring this work to fruition: Dr Anna Laddach, who developed the original TITINdb site and gave much of her time for troubleshooting in the initial updates; Dr Martin Rees, who consistently gave helpful feedback to us about the performance and features; Rob Wilson and Denis Mills, who worked with us to migrate the TITINdb2 server and deliver updates.

