## Supplementary Material for "TITINdb2 – Expanding Annotation and Structural Information for Protein Variants in the Giant Sarcomeric Protein Titin"

### Application Features

We here list the intended features for which TITINdb2 was developed, and are available in the current version of the database:

- A curated database of missense single amino acid variants (SAVs) in titin, aligned to the largest continuous isoform of titin, the inferred complete (IC) isoform (Ensembl Transcript ID: ENST00000589042.5 (Martin *et al.* 2023); NCBI Reference Sequence: NM_001267550.2 (O’Leary *et al.* 2016)).
- Access to all missense SAVs reported in the population SNV databases 1KGP (The 1000 Genomes Project Consortium 2015), and the Genome Aggregation Database (gnomAD) version 2.1.1, version 3.1.2, and version 4.0.0 (Karczewski *et al.* 2020), including population minor allele frequencies, and sub-population allele frequencies where available.
- Access to reported disease-associated variants in the literature, as well as links to references in PubMed and a short summary of available evidence.
- Population and disease-associated variants are annotated with pathogenicity prediction scores through cross-reference with large-scale prediction databases dbNSFPv4 (Liu *et al.* 2020b), FannsDB (FannsDB) and can be accessed through the "Custom Data Table" dialog option. These predictors include AlphaMissense (Cheng *et al.* 2023a), Condel (González-Pérez and López-Bigas 2011), REVEL (Ioannidis *et al.* 2016)), and MVP (Qi *et al.* 2021); for a full list of predictors in TITINdb2 see Supplementary Table 2.
- Population and disease-associated variants are further annotated with protein stability prediction scores calculated using DUET (Pires, Ascher and Blundell 2014a) and functional site annotations from UniProt.
- All variant information and annotations can be downloaded for a single domain as a single CSV file. Protein sequences for single domains can be downloaded as a single file in FASTA format. Data tables for titin are made easily available for download in CSV or JSON format, as well as bulk data downloads in compressed formats.
- Access to new and emerging data from AI and large language model-based approaches: single- and tandem-domain structural models of titin predicted by AlphaFold2 (Jumper *et al.* 2021), extracted from proteome predictions in AlphaFoldDB (AlphaFold Protein Structure Database); predicted pathogenicity scores for all possible SAVs at every position in titin for both the IC and canonical (N2AB) isoforms predicted by AlphaMissense (Cheng *et al.* 2023a), extracted from proteome predictions distributed via Zenodo (Cheng *et al.* 2023b).
- All possible SAVs in all titin domains (IC isoform) are annotated with protein stability predictions calculated using mCSM (Pires, Ascher and Blundell 2014b), as well as all possible SAVs in titin domains (N2AB isoform) annotated with pathogenicity predictions calculated using Rhapsody (Ponzoni *et al.* 2020) and PolyPhen-2 (Adzhubei, Jordan and Sunyaev 2013), available through the saturation mutagenesis feature.
- Mapping of SAVs to experimental structures from PDB (see Supplementary Table 2), as well as homology models predicted by Modeller (Eswar *et al.* 2006), and an ensemble of models predicted by AlphaFold2 (Jumper *et al.* 2021; AlphaFold Protein Structure Database). All structures are available for download as PDB files, as well as the 1400-residue AlphaFold2 "contig" predictions.
- Relative quality of structures and models for individual or tandem domains can be compared through experimental resolution for X-ray crystal and electron microscopy structures, and MolProbity (Williams *et al.* 2018) score and Rama Z-score (Sobolev *et al.* 2020) for models as an estimate of effective resolution.
- SAVs localising to structures are annotated with structure-specific analysis: residue quotient solvent-accessible surface area (Q(SASA), computed with POPS (Cavallo, Kleinjung and Fraternali 2003)); predicted protein-protein interaction residues (computed with SPPIDER (Porollo and Meller 2007)).
- Visualisation of individual SAVs on the corresponding domain structure; images can be saved as PNG files.
- Visualisation of distribution of SAVs across a given domain on the corresponding domain structure, from the databases listed above.
- Visualisation of domain-domain interface residues as determined using AlphaFold2 predicted single and tandem domains.
- Search for variants by isoform-specific amino-acid position, by dbSNP rsID, or by the associated disease or pathology.
- Comprehensive re-mapping of amino acid positions to transcript reference sequences, chromosomal reference sequences; these data are available through the "Position" page on TITINdb2. Variants may also be converted between protein, transcript, and genomic positions in different isoforms using the "variant converter" tool available through the web server.

### Case Studies and Applications

#### Comparison of Variants with Pathogenicity Prediction Methods

Next-generation sequencing (NGS) and other high-throughput approaches generate huge quantities of data, including the genetic data from patients with inherited diseases. In the case of titin, not only have variants in titin been associated with a wide range of muscle diseases, but the sheer size of titin also means that all individuals will contain some missense variants of varying significance within their titin sequence. By extension, this also means that healthy individuals will have missense variants, often those which would be considered rare in the general population. This complicates the prioritisation of variants, where the great quantity of variants identified in a patient population must be filtered to produce a candidate list of a size amenable to further experimental validation.

Bioinformatics annotation pipelines are often used to identify variants of interest from within large gene sequencing data sets, by annotating the individual variants with bioinformatics tools and then filtering by specific criteria. TITINdb2 can perform a similar function for titin missense variants, but with certain advantages over these more general approaches; like other annotation pipelines, it provides minor allele frequency (MAF) values for the population variant databases, as well as those of reported sub-populations, but it also provides all population and computational information in a single location, without the need for separate software or converting between variant formats. For example, the availability of pre-computed pathogenicity predictions for variants of interest means that the user need not run these programs for each variant individually.

As an example, we will illustrate the use of TITINdb2's new Custom Data Table dialogue to produce relevant annotations on variants. The Ig-169 domain of titin is considered a hotspot for variants associated with tibial muscular dystrophy (TMD) and limb-girdle muscular dystrophy type 2J (LGMD2J) (Hackman *et al.* 2002), with multiple missense and insertion-deletion variants reported associated with these conditions. Moreover, a study that characterised the effects of these variants on the Ig-169 domain *in vitro* reported a measurable spectrum of severity between different variants, based on assays of stability, solubility, expression, and binding to OBSCN Ig-1 (Rudloff, Woosley and Wright 2015). The three missense variants thus characterised, L35965P(Hackman *et al.* 2002), H35946P (Pollazzon *et al.* 2010), and I35947N (Van den Bergh *et al.* 2003), are also distinguishable in other respects; while L35965P and H35946P are fully penetrant and absent from population databases, I35947N, the variant characterised as least severely damaging, is also found at low frequency in population databases (see Supplementary Table 3), was found with incomplete penetrance in the source population, and is thought to be only pathogenic when homozygous or compound heterozygous with other variants (e.g. Q22507* (Evilä *et al.* 2017)).

Since the characterisation of these variants, two further missense variants associated with disease in Ig-169 have been identified by genetic analysis of genes in affected families: W35930R (LGMD) (Zheng *et al.* 2016; Evilä *et al.* 2017) and T35915P (TMD) (Evilä *et al.* 2017). As with the previous variants, there are differences in the penetrance displayed; W35930R co-segregates fully with the disease in the 6 individuals tested (3 carriers, 3 non-carriers), whereas T35915P co-segregates with disease only when compound heterozygous with the frameshift variant E28338fs, like with I35947N. While no experimental characterisation on these variants has yet been performed, we can use TITINdb2 to compare these variants *in silico* using pathogenicity prediction annotations.

Using the Custom Data Table, we select only those pathogenicity predictors that were published and last updated before 2016, when the two newer missense variants were identified; we also include AlphaMissense, which while published in 2023 does not incorporate pathogenic/neutral labels in its learning algorithm. While this excludes most predictors from the data set, it also demonstrates how one can select only those predictors desired when performing a task that requires only a specific subset of the data available, and how TITINdb2 facilitates this (for a full list of predictors in TITINdb2 and the date of their publishing, see Supplementary Table 2). The predicted values for each of the five variants may be seen below:

Supplementary Table 1: Predicted pathogenicity scores for damaging variants located in titin's Ig-169 domain. Prediction scores differ on scale and weighting; variant scores are coloured orange for higher likelihood of the variant being pathogenic, and green for lower. AlphaMissense, MutationTaster, PolyPhen-2 and REVEL are bounded by values of 0 to 1; other predictors do not have minimum or maximum scores.

| **Variant** | **Alpha-Missense (IC)** | **FATHMM** | **MetaLR** | **MetaSVM** | **Mutation-Assessor** | **Mutation-Taster** | **PolyPhen-2** | **PROVEAN** | **REVEL** |
| --- | --- | --- | --- | --- | --- | --- | --- | --- | --- |
| **L35956P** | 1 | -0.75 | 0.68 | 0.55 | 3.6 | 1 | 1 | -5.02 | 0.79 |
| **H35946P** | 0.78 | -0.24 | 0.41 | -0.2 | 1.9 | 1 | 1 | -6.46 | 0.63 |
| **I35947N** | 0.73 | -0.45 | 0.42 | 0.14 | 2.95 | 0.83 | 0.9 | -4.87 | 0.65 |
| **W35930R** | 1 | -3.97 | 0.97 | 1.09 | 4.55 | 1 | 1 | -9.87 | 0.94 |
| **T35915P** | 0.79 | 0.79 | 0.37 | -0.12 | 3.26 | 0.99 | 1 | -3.97 | 0.47 |

As can be seen in Supplementary Table 1, the predicted pathogenicity scores for the three characterised variants correlate well with their known severity; L35956P is consistently calculated as the most likely damaging of the three, and has the maximum pathogenicity score from AlphaMissense, MutationTaster and PolyPhen-2. I35947N is typically calculated as the least damaging variant, and never the most damaging, while H35946P is more variable between predictors, averaging to an intermediate value. Similarly, W35930R is computed as severely damaging, and matches or even exceeds the value of L35956P In all predictors; by contrast, T35015P shares the inconsistent prediction of H35946P, averaging to intermediate between H35946P and I35947N. From these predictions, we might hypothesise that their effects if characterised *in vitro*, as the first three variants were, would fall into similar places on the spectrum.

#### Congenital myopathy-associated variants in the A-band

The concept of the "hotspot", as it relates to titin, refers to a position or positions in a molecule that are highly intolerant to mutation – thus, variants at these positions are typically associated with damaging consequences. As titin is a large, modular protein, the identification of its hotspots is somewhat less intuitive – variants in different regions of the protein may have greatly differing effects, potentially realised only in specific tissues. Thus, hotspots in titin can be categorised in two ways: firstly, hotspot domains, where many damaging variants within a single domain or closely situated domains result in a similar phenotype; secondly, hotspot positions, wherein many damaging variants may be located at or around a similar structural location in different domains with the same fold. In the previous version of TITINdb, a case study demonstrating how these hotspots may be visualised on structural models of domains was provided; here, we expand on this analysis to highlight new features in TITINdb2 that allow even further characterisation of these positions.

It is well-established in the literature that variants in the Fn3-119 domain of titin are strongly associated with hereditary myopathy with early respiratory failure (HMERF) (Palmio *et al.* 2019), with 13 reported missense variants in the literature alongside the condition. Many of these variants occur at the same location in the domain; for example, C31712R, C31712Y and C31712W all occur at position 5 in Fn3-119. This suggests that within the fibronectin domain fold are specific residues integral to the stability of the domain and poorly tolerant of mutations; this observation has been reinforced with the identification of disease-associated variants in other domains that closely mirror those already identified in Fn3-119 (Rees *et al.* 2021). In four of these cases, A-band variants identified in patients with multi-minicore disease (MmD) represented the exact same residue change, at the exact same structural location, as variants reported in HMERF patients in Fn3-119 (see Supplementary Figure 1).

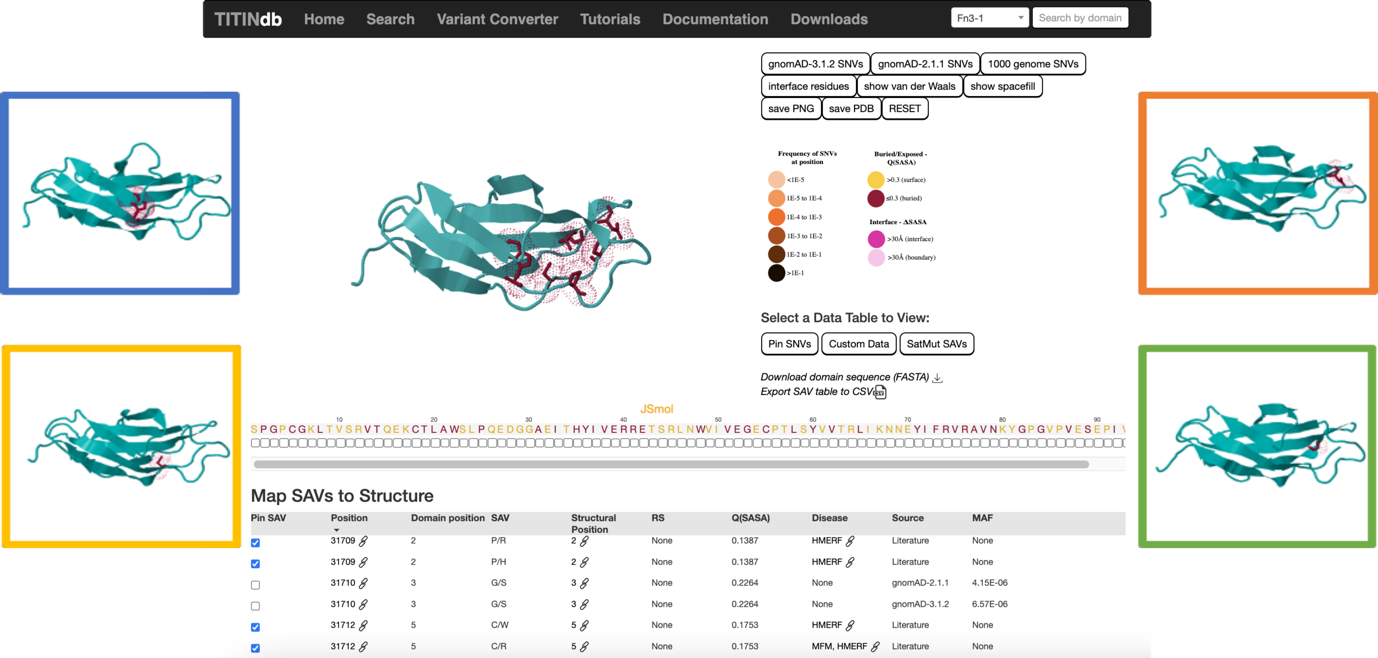

Supplementary Figure 1 – A screen capture of TITINdb2 web server querying the Fn3-119 domain, using a predicted structural model sourced from AlphaFold2. All known disease-associated variants within this single module have been "pinned" to visualise their respective structural locations. A simple colour code of maroon for these residues illustrates that these are all located in buried, mostly solvent-inaccessible positions, versus yellow for surface residues. Inset, disease-causing variants in other titin Fn3-type domains are located at the same structural positions as those in Fn3-119 and represent the same amino acid substitution: blue, W16471C in Fn3-7 mirrors W31729C in Fn3-119; orange, N16133K in Fn3-4 mirrors N31786K in Fn3-119; green, G27849V in Fn3-90 mirrors G31791V in Fn3-119; yellow, C17051R in Fn3-12 mirrors C31712R in Fn3-119.

As in the previous version of TITINdb2, by navigating to the page for the specific domain of interest and selecting the variants, or residue positions, of interest, the user may visualise the position of those residues on a representative domain structure. Here, we can verify that the positions of these variants in their respective domains match; this reinforces the idea that certain positions in these domain types are more susceptible to pathogenic variants than others. In the newest version of TITINdb2, we can additionally search for these variants at shared structural positions directly, using the "Structural Position" annotation. By navigating to either the domain or position page in which a variant of interest is located, the "Structural Position" annotation in the SAV table links through to a new page in which all variants located at that aligned structural position in all homologous domains in titin may be accessed.

The wild-type asparagine residue that is mutated in the N31786K (Fn3-119) and N16133K (Fn3-4) variants is highly conserved across titin Fn3 domains, and by navigating to its structural position page, we can see that it is typically located at domain sequence position 79-85, depending on the domain in question. Interestingly, the structural position page identifies two other disease-associated ASN substitutions in other domains at this same structural position: The AVB-associated N16429K variant (Fn3-6) (Liu *et al.* 2020a) and the congenital titinopathy-linked N19955I variant (Fn3-32) (Yu *et al.* 2019). Both variants have yet to be characterised experimentally to conclusively demonstrate their pathogenicity, but their disruption of the same position as two other known damaging variants suggests that these bear closer examination. However, the structural position page also highlights that there are ASN-to-LYS substitutions at this same position reported in 15 other Fn3 domains, in individuals in the general population, and thus that the penetrance and severity of these variants may be variable.

#### Distributions of Saturation Mutagenesis Predictions in TITINdb2 Populations

Saturation mutagenesis, or deep mutational scanning (DMS), refers to the systematic evaluation of the impacts of all possible single protein residue alterations at all positions; in experimental protocols, this involves high-throughput assays assessing a measurable parameter for protein constructs, such as stability or binding affinity (Fowler and Fields 2014). Similarly, computational mutational scans using pathogenicity predictors allow for comparison of vast numbers of protein substitutions in large proteins such as titin. TITINdb2 incorporates saturation mutagenesis data from several different prediction tools; three such tools are AlphaMissense, Rhapsody and REVEL. AlphaMissense (Cheng *et al.* 2023a) is a recently released large language model-based algorithm based on the AlphaFold2 architecture; Rhapsody is a relatively recent algorithm that incorporates potential pathogenicity-associated features of variants from a variety of data types; REVEL is a meta-predictor that incorporates a consensus approach to render a final pathogenicity score.

With the release of AlphaMissense and the success of AlphaFold2 in achieving the first highly accurate *ab initio* protein structure prediction tool based on deep learning methods, much work is currently in progress to determine if AlphaMissense represents a similar advance in the field of variant impact prediction in the context of different disease-associated proteins. For titin, a gene and protein in which the number of potential variants vastly exceeds the practical means for any current method to experimentally characterise them, this is particularly important; therefore, we obtained the annotations for AlphaMissense, Rhapsody, and REVEL for variants reported in the different population variant databases in TITINdb2, as well as those for variants reported pathogenic in the literature with experimental validation, and compared the distributions of predictions across databases and across methods.

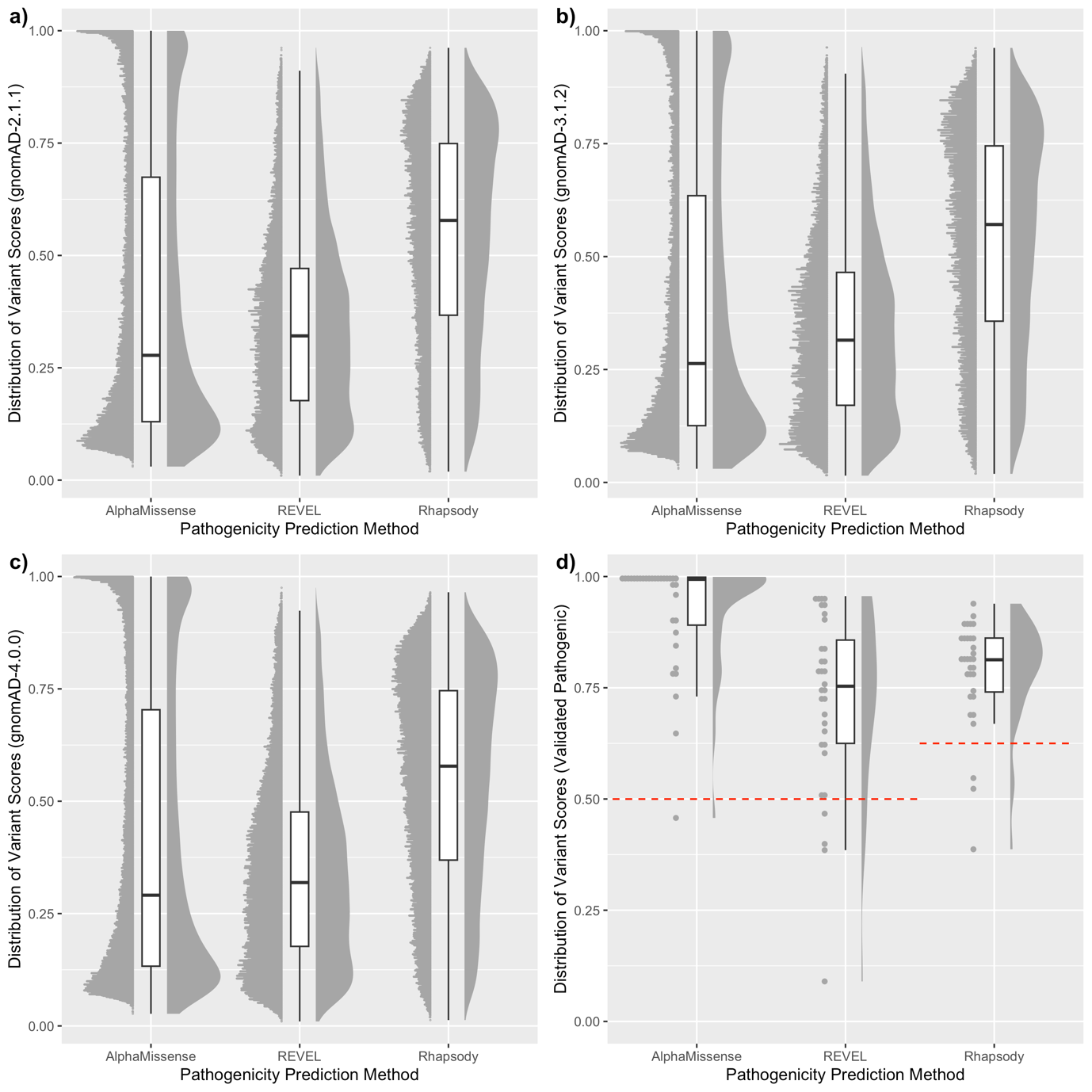

Supplementary Figure 2 – Distributions of predicted pathogenicity scores computed by a) AlphaMissense (IC Transcript), b) Rhapsody, c) REVEL. All 3 algorithms output a score between 0 and 1, where 1 indicates the highest possible certainty of the pathogenicity of the variant, and 0 the lowest. Four populations of SAVs are represented here – three databases of population-derived SAVs (gnomAD-2.1.1, gnomAD-3.1.2 and gnomAD-4.0.0) and variants reported pathogenic in the literature with substantial evidence in support of their pathogenicity. Each population is visualised by three separate plots to illustrate the distributions of predictions: on the left, a histogram plot representing the binned frequencies of predictions; in the middle, a box plot representing range, median, and interquartile range of the distribution of predictions; on the right, a violin plot representing the relative distributions of predictions. For variants reported pathogenic, a red line indicates the typical threshold value used to classify as pathogenic or neutral (0.5 for AlphaMissense/REVEL, 0.625 for Rhapsody).

As seen in Supplementary Figure 2d, all three predictors predict the majority of the 31 validated pathogenic variants reported in the scientific literature correctly, with AlphaMissense correctly predicting all but one of these variants. However, the shape of the distributions across population databases (Supplementary Figure 2a-c) differs between the algorithms. The distribution of AlphaMissense predictions is bimodal, with the most populous bins by frequency representing those variants with extremely high or extremely low likelihoods of pathogenicity. By contrast, the distribution of Rhapsody predictions is more balanced, with these extreme values representing the least populous bins in the databases, and predictions more evenly spread over the intermediate bins of the distribution. Finally, the distribution of REVEL predictions shows a skew towards predicted values below 0.5, or pathogenicity scores that would classify the variant as neutral.

This comparison is illuminating in several ways, both in terms of how the algorithms approach the question of pathogenicity and how they may be best used in practical circumstances. AlphaMissense tends to make predictions with high certainty, with variants either very likely pathogenic or very likely neutral, as per its goal of binary classification of variants into these two categories. Rhapsody, by contrast, makes predictions with a more pronounced spectrum of degree of pathogenicity, as per the number of different features that may argue for or against the damaging quality of the variant. Finally, REVEL predicts most variants in the population data sets as non-pathogenic, which intuitively should be the case.

Because of these differences, each of these methodologies may be better suited to certain applications than others. For example, the bimodal classification of AlphaMissense makes it less sensitive to classification thresholds, and thus may be better suited to variant prioritisation tasks where variants are retained or discarded from consideration based on computational characterisation. Rhapsody, which shows a greater degree of separation between variants considered pathogenic or non-pathogenic, may be better suited to investigating the differences between the severity or penetrance of different pathogenic variants in the same system. Finally, REVEL, which appears to predict fewer false positives than the other methods, may be best applied to tasks involving the processing of background mutational noise.

### Methods

#### Data Collection

Population missense variant data is sourced from four databases: the 1000 Genomes Project (1KGP)(The 1000 Genomes Project Consortium 2015); gnomAD version 2.1.1, gnomAD version 3.1.2, and gnomAD version 4.0.0 (Karczewski *et al.* 2020). Missense variants in the 1KGP data set were retrieved using the online data slicer (The 1000 Genomes Project Consortium 2015) and mapped to the inferred complete isoform; these originate from 2504 nominally healthy individuals. The gnomAD data sets are aggregate databases comprising many large-scale genome sequencing studies; the v2.1 data set includes data from 125,748 exomes and 15,708 genomes from 141,456 unrelated individuals, while the v3.1 data set includes data from 76,156 genomes from unrelated individuals. The gnomAD database v4.0 is composed of 730,947 exomes and 76,215 genomes and is reported to include all data from the 2.1.1 and 3.1.2 data sets; however, there are a non-negligible number of variants in titin present in one or the other of these older releases that are not reported in the newest database. Additionally, v4.0 possesses a much higher sampling of individuals with European ancestry compared to the older releases, thanks to the inclusion of data from the UK Biobank (Van Hout *et al.* 2020); thus, minor allele frequency (MAF) and sub-population frequencies for all three databases are given in separate data entries within TITINdb2.

While individuals with severe paediatric disease and their close relatives are excluded from gnomAD, the data sets also include disease-specific genetic studies, so their inclusion in the population data does not necessarily preclude pathogenicity. However, in general the gnomAD data set should not be enriched for damaging variants. In addition, while all three databases give sub-population allele frequencies, the ethnic populations considered for each database differ; most notably, the sub-population "European" is given for 1KGP variants, whereas all gnomAD versions split this sub-population into "Finnish" and "Non-Finnish European". Further information can be found on the online Documentation page.

Reference SNP cluster IDs (rsIDs) matching protein variants found in TITINdb2 were previously sourced directly from gnomAD annotations; however, the mapping of rsIDs to protein variants here is incomplete. Thus, we downloaded records for all rsIDs for missense variants in titin from dbSNP (Sherry *et al.* 2001), then converted the given chromosomal coordinates to the corresponding protein coordinates using the variant converter tool (see below; "Mapping of Protein, Transcript and Chromosomal Data"). Protein variants in the database were then annotated with the matching rsID; note that, as rsIDs include all reported nucleotide substitutions at that position, one rsID may correspond to more than one protein variant.

#### Extraction of Predicted Domain Structures and Variant Annotations from AlphaFold and AlphaMissense

The coverage of experimentally solved titin domains in the literature has continued to expand, with 48 entries in the PDB corresponding to domains in titin, covering 33 domains in total (see Supplementary Table 3). However, there are still 268 domains in titin that lack structural coverage. In the original build of TITINdb, a computational pipeline was constructed to define the domain boundaries for all detected domains in titin's inferred complete (IC) isoform, and then to model those domains using the template-based modelling software, Modeller (Eswar *et al.* 2006). In brief, domains were defined in the IC isoform by scanning with HMMER (Finn, Clements and Eddy 2011) against Pfam seed libraries (Finn *et al.* 2014) and validated by generation of sequence logos (Weblogo (Crooks *et al.* 2004)) from multiple sequence alignments of Ig/Fn3 domains thus defined (T-Coffee (Notredame, Higgins and Heringa 2000)) and alignment with existing experimental structures of homologous titin domains. These individual domain sequences in FASTA format were then supplied to Modeller as input for template search, homology modelling, and model assessment steps. The full pipeline, including schematic figure, can be found in the original paper.

Recent advances in the field of protein structure prediction have culminated in the release of AlphaFold2, a Deep Learning-based software capable of *in silico* predictions of protein structural models from sequence data alone (Jumper *et al.* 2021). Moreover, the authors have released the protein structure predictions for all proteins in the proteomes of many species via the online database AlphaFoldDB (AlphaFold Protein Structure Database). However, the size of titin renders any attempt to model the protein in its entirety impractical; thus, predictions for titin domains are not readily accessible through standard searches on the AlphaFoldDB web portal, and users must navigate the large proteome database to extract pre-computed titin structure predictions.

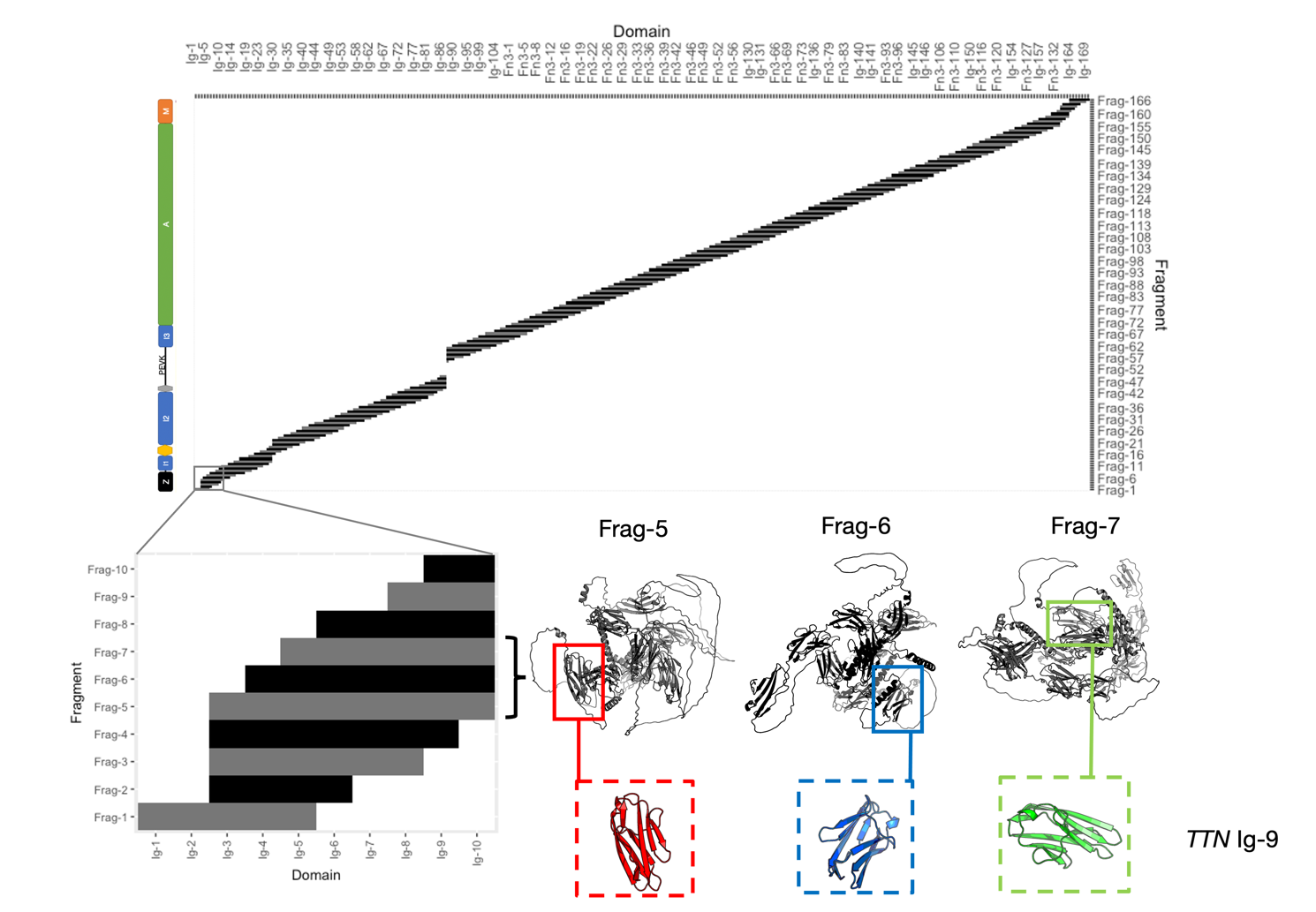

Supplementary Figure 3 - Overview of the extraction of AF2 domain structures from 1400-residue fragments. Each fragment is searched for a complete match to the sequence of each domain in titin, and matching residues are extracted to a new PDB object. For example, the domain Ig-9 is found in fragments 5-11, each of which produces a slightly different domain structure prediction.

To improve the usability of these data for the study of titin, we have processed and extracted these domain systems, and they are available via the TITINdb2 web portal. The full proteome predictions for *H. sapiens* were downloaded from AlphaFoldDB and the entries corresponding to the UniProt entry for titin's N2AB isoform were extracted. These are organised into what we denote "contigs" of overlapping predictions; a sliding window of 1400 residues is taken from titin and used to compute a predicted structure, moving in "steps" of 200 residues. Thus, the whole titin N2AB sequence is represented in 166 contigs, with each individual domain represented fully in 6-7 contigs. The coverage of the 302 domains in titin across the 166 contigs is visualised in Supplementary Figure 3; the structures extracted reinforce the previous assessment of locations of domains in titin.

A pipeline was developed to identify and extract all single and tandem domains from these contigs, using the previously established domain boundaries. Tandem domains were defined as successive domains separated, according to the domain boundaries, by a linker of 10 or fewer residues. The pipeline, written in Python 3.8 with the use of the PyMol module (Schrödinger, LLC 2015), took the sequences for the single and tandem domains in FASTA format as input and searched for a sequence match within the sequences for the supplied contigs; if a successful match was found, that segment of the contig was extracted and saved as a new PDB file. The pipeline produced a total of 1925 single-domain and 1624 tandem-domain structures. As each of the models for a given titin domain were produced in a slightly different chain context, there are differences between them; this is especially pronounced for tandem domains with long linkers. As such, the web server provides for visualisation of variants on all the single and tandem domains extracted from contigs, as well as the capability to download the original contigs from the structure selection page.

As the UniProt canonical sequence was used by AlphaFold2 to model all contigs, there is no coverage of the novel exon domains Ig-25 or Ig-26, nor the 6 novex-3 domains found in exon 48 (Ig-25_novex3 to Ig-30_novex3). To remedy this, we used ColabFold (Mirdita *et al.* 2022), which incorporates the slightly less extensive but much faster multiple sequence alignment tool MMseqs2 (Steinegger and Söding 2017), rather than the jackhmmer (Potter *et al.* 2018) algorithm used by AlphaFold2. Five models for each domain were produced by the ColabFold notebook, from which the best-ranked model by pLDDT was selected and included in TITINdb2.

#### Validation and Comparison of Model Structures

In order to compare the AlphaFold2-derived models to the previously generated homology models and PDB experimental structures, we calculated two measures of model quality using a local version of the MolProbity (Williams *et al.* 2018) pipeline in Python and Phenix. The first of these, the MolProbity score, is an estimate of the effective resolution of the model were it an experimentally solved structure; it incorporates the MolProbity clash score with Ramachandran and rotamer outliers to give a single value which reflects the experimental resolution at which such a score would be on average. Thus, the closer the value to zero, the better the assessed quality of the model. The second measure, Ramachandran Z-score (Sobolev *et al.* 2020), summarises the Ramachandran outliers/favoured in a single value, where $abs(Z)\leq2$ would generally be considered to represent a good quality model. The MolProbity score was calculated for all AlphaFold2 and Modeller models and compared against the actual experimental resolution of X-ray crystal structures in the literature.

The single domain AlphaFold2 models compared very favourably to the homology models and experimental structures, with the best models approaching an estimated resolution of around 0.5 Å, and all models under 2.5 Å (see Supplementary Figure 4a). Similarly, the AlphaFold2 models are considered very good models based on the Rama-Z score, although here the homology models are comparable in quality to the average AlphaFold2 model, and only significantly lower quality than the AlphaFold2 models judged best (see Supplementary Figure 4b). For the ColabFold models, the assessed quality is of a similar level to that of the AlphaFold2 models, with MolProbity scores of 1.57 (Ig-25) and 1.70 (Ig-26). Overall, the provision of AlphaFold2 models has increased the quality of the structural coverage for titin.

We caution the reader that Ig-112 remains a problematic domain for modelling. This domain contains a large insert within a loop region, which is poorly modelled by all methods; in the previous build of TITINdb2, it could not be modelled to an acceptable standard using Modeller, and as such was modelled with I-TASSER (Zhang 2008). The AlphaFold2 models satisfy geometric constraints adequately, and do not appear as outliers in terms of their MolProbity and Rama-Z scores; however, be aware that its prediction for the loop does not reflect any existing experimental knowledge.

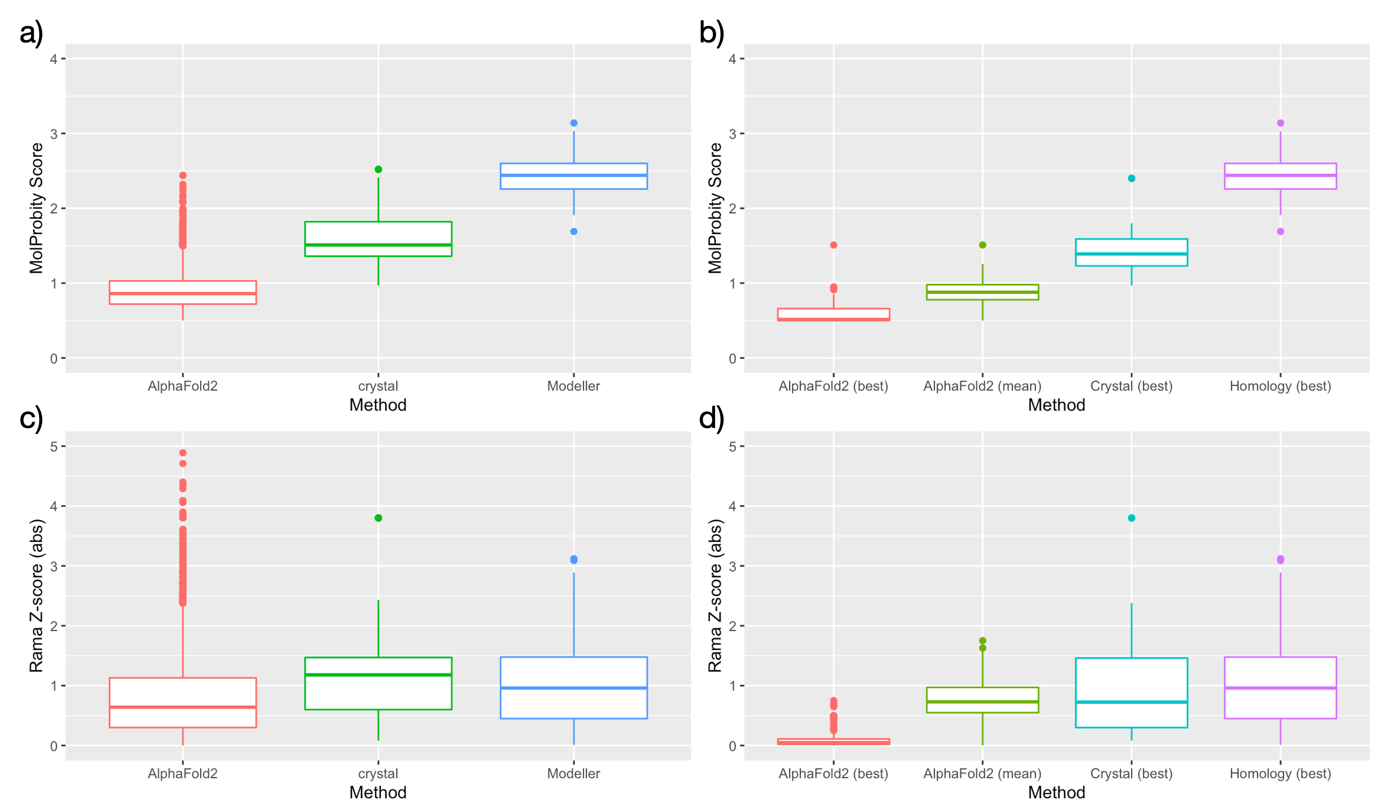

Supplementary Figure 4 – Comparisons of estimated structural quality between predicted and experimentally solved structures using different methods. AlphaFold2-derived models are estimated to be of high accuracy both by MolProbity score (a, b) and Rama Z-score (c, d). Both best and average AlphaFold2 models per domain are significantly better than the best crystal and homology models (b), but there is no difference in quality according to Rama Z-score (c, d).

#### Annotation of Missense Variants

A subset of known variants are annotated with 13 pathogenicity prediction scores sourced from the annotation aggregation database dbNSFPv4 (Liu *et al.* 2020b); a full list of these predictors and their original publication can be found in Supplementary Table 2. TITINdb2 also includes saturation mutagenesis predictions for all possible variants in titin, both reported and theoretical: pathogenicity predictions for all possible nsSNVs in titin's canonical isoform are available for the meta-predictors Condel (González-Pérez and López-Bigas 2011) and REVEL (Ioannidis *et al.* 2016); Condel predictions were obtained from FannsDB (FannsDB), while REVEL predictions were sourced from their genome segment database (REVEL genome segment files) and mapped from chromosomal to protein coordinates before being added to TITINdb2.

Saturation mutagenesis pathogenicity predictions for all SAVs present in titin's canonical (N2AB) isoform were extracted from the pre-computed databases for PolyPhen-2 (Adzhubei, Jordan and Sunyaev 2013) using their online web server. These predictions were then used as input for saturation mutagenesis predictions for all N2AB SAVs located within domain boundaries for Rhapsody (Ponzoni *et al.* 2020), a predictor which uses sequence-, structural-, and dynamics-based features to predict pathogenicity. As Rhapsody's web server does not have access to the full structural coverage of titin available in TITINdb2, we implemented a local pipeline incorporating pre-computed PolyPhen-2 predictions and titin domain models from AlphaFold2 (see below) to generate predictions.

As in the previous build of TITINdb2, all possible SAVs within domain boundaries in the inferred complete (IC) isoform of titin were assessed using mCSM (Pires, Ascher and Blundell 2014b) to predict protein stability; these were re-calculated using AlphaFold2 models. The previous protein stability predictions made by DUET (Pires, Ascher and Blundell 2014a), for all possible nsSNVs only, are also included. Finally, the recent release of AlphaMissense (Cheng *et al.* 2023a) has included pre-computed predictions for all possible SAVs at every position in titin, both for the canonical N2AB isoform as given by UniProt (ID QW8Z42), and for the inferred complete isoform as given by Ensembl (Martin *et al.* 2023) (transcript ID ENST00000589042.5). As these predictions contain minor differences, both are included in saturation mutagenesis data in TITINdb2 for the sake of comparison; both are sourced from the distribution of AlphaMissense predictions through Zenodo (Cheng *et al.* 2023b).

Variants in TITINdb2 are annotated for predicted protein-protein interactions using SPPIDER (Porollo and Meller 2007), which is expressed as a binary classification; variants in domains where an experimental structure exists for that domain in complex with an interaction partner (PDB 1YA5 for Ig-1/Ig-2 in complex with TCAP; PDB 3KNB for Ig-169 in complex with OBSL1; PDB 4UOW for Ig-169 in complex with OBSL1; PDB 4C4K for Ig-169 in complex with OBSCN) have been assessed with mCSM-PPI (Pires, Ascher and Blundell 2014b) to predict the impact of that mutation on the binding affinity between the two proteins.

#### Definition of Structural Regions of Domains

In the previous build of TITINdb2, the quotient solvent-accessible surface area – Q(SASA) – was calculated for all residues in all domains of titin and used to annotate the variants in the database. This was performed automatically by the tool POPS (Cavallo, Kleinjung and Fraternali 2003), using the default probe radius of 1.4Å. Residues with $Q(SASA)>0.3$ were defined as exposed/surface residues, while those with $Q\left( SASA \right)\leq0.3$ were defined as buried/core residues. Additionally, with the availability of tandem titin domains through AlphaFold2, we calculated interface regions for all tandems defined according to the previous boundaries.

$$if {Q(SASA)}_{single}>0.3 then "EXPOSED"$$

$${if Q\left( SASA \right)}_{single}\leq0.3 then BURIED"$$

$$if {SASA}_{single}-{SASA}_{tandem}\geq30Å^{2} then "INTERFACE"$$

For each residue in the tandem, the SASA value of the residue in the tandem domain model was subtracted from the SASA value of that same residue in the single domain to give the change in SASA, or ΔSASA, an adaptation of the method used in POPSCOMP (Kleinjung and Fraternali 2005) for protein complexes. Thus, a value of $\Delta SASA\geq30 Å^{2}$ for a particular residue would be considered an interface-buried residue. An example of this workflow is provided in Supplementary Figure 5.

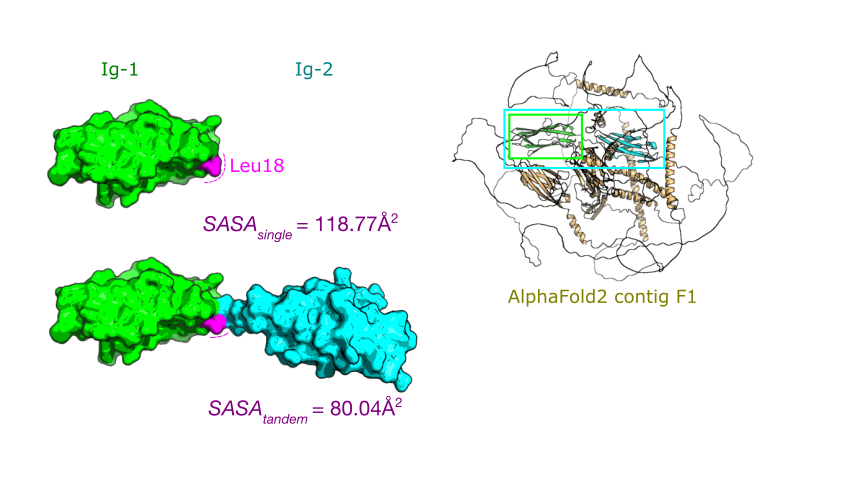

Supplementary Figure 5 - Example interface-buried region in titin Ig-1. Solvent-accessible surface area (SASA) of the leucine residue at position 18 is decreased by 38.73 Å^2^ when Ig-2 is included in the construct, thus this residue is annotated as interface-buried. Right, AlphaFold2 structure F1, from which the structures for Ig-1 and Ig-2 used here are derived.

#### Mapping of Protein, Transcript, and Chromosomal Data

To produce the Variant Converter tool that would enable users to translate the coordinates of protein, transcript, or chromosomal variants to any other format, we conducted a mapping of reference sequences of each type to one another, using the seven major isoforms of titin (IC, N2AB, N2B, N2A, novex-1, novex-2, novex-3). Genomic sequences and positional data for the hg19 and hg38 builds of the human reference genome were sourced from CardioDB (Roberts *et al.* 2015; Titin Variation in Dilated Cardiomyopathy - Cardiovascular Genetics & Genomics, Imperial College London), transcript and protein sequences corresponding to the seven isoforms were sourced from RefSeq (O’Leary *et al.* 2016). The data were incorporated into TITINdb2 as annotations for each IC isoform position; these included genomic and transcript codons, corresponding positions in transcript for all isoforms (e.g. "758,759,760") and corresponding positions in both hg38 and hg19 genome for all isoforms (e.g. "178800445,178800444,178800443").

The Variant Converter tool pulls from these annotations to translate variants quickly and reliably in bulk between protein, transcript, and chromosomal positions. The conversion from SAV to SNV relies on coercion of the wild-type codon to a mutant codon matching the mutated residue; if multiple codon substitutions match the amino acid substitution, the tool will only report the first match. Additionally, the tool only supports the conversion of SNVs and SAVs; it will not, for example, translate an SAV that corresponds to an indel at the genetic level (e.g. the SAV C31712R may be coerced to the SNV 95359T>C, but the theoretical SAV C31712K could not be coerced to the indel 95359_95360delinsAA). The Variant Converter does not recognise the standard HGVS formatting for variants, but rather uses the simple variant nomenclature syntax outlined below:

| **Format (IC)** | **Example HGVS Nomenclature** | **TITINdb2 Variant Converter** |
| --- | --- | --- |
| Chromosome | NC_000002.12:g.178546102A>G | 178546102A>G |
| Transcript | NM_001267550.2:c.95359T>C | 95359T>C |
| Protein | NP_001254479:p.Cys31712Arg | C31712R |

In addition to selecting isoform and format by name, users may select input and output formats using accession IDs derived from RefSeq, UniProt and Ensembl. This tool was used to convert chromosomal positions for SNVs in titin to their translated protein notation for REVEL predicted pathogenicity scores, as well as for reference SNP records from dbSNP.

Two possible sources of confusion are addressed here. First, the 225bp upstream non-translated sequence is included in the numbering for all titin transcripts; thus, the ATG start codon for the initial methionine residue in all isoforms is numbered as 226-228. This may be inconsistent with other numbering schemes in the literature that omit these nucleotides, thus users should be aware of these potential differences. Second, certain codons in titin are split across exons, meaning that genomic numbering is not always sequential; furthermore, because of exon usage, it is possible for a chromosomal variant that produces a certain protein variant in one isoform to produce a wholly different one in another isoform. For example, the variant G3435R in the IC isoform results from a single nucleotide substitution that instead produces the variant A3389P in the N2B isoform.

#### Structural Positions and Alignment

While there are limited numbers of variants reported at each individual position in titin, the Ig and Fn3 domains that make up the protein have a high sequence identity, and information about variation at a given position in a domain may be provided by variants at the same structural position in homologous domains. Thus, in the new version of TITINdb2 we defined structural positions in titin according to shared positions in the topology of the domain fold, independent from the absolute domain sequence position. Aligned structural positions were allocated based on multiple sequence alignments of all 169 Ig domains and 132 Fn3 domains in titin; the structural position annotation, accessible as a link from the page for any residue located within a domain, allows the user to navigate to a page where all reported variants at that shared structural position, as well as the wild-type residue and the domains they occur in, can be viewed.

Consensus structural alignments were generated using PROMALS3D (Pei, Kim and Grishin 2008) with available high-quality experimental X-ray crystal structures from PDBe for both Ig and Fn3 domains; these were then mapped back onto the multiple sequence alignments of all Ig and Fn3 domains, respectively, and shared structural positions were annotated with consensus secondary structure information at that position.

### Supplementary Tables

*Supplementary Table 1 can be found above, Supplementary Tables 2, 3 and 4 below.*

##### Supplementary Table 2

*Predictors used to annotate variants in TITINdb2. A brief description of the methodology and a reference to the original paper is given, as well as the coverage of annotations across titin variants.*

| Predictor Name | Source | Description | Availability | Reference |
| --- | --- | --- | --- | --- |
| AlphaMissense | Predictions for AlphaMissense (Zenodo) (Cheng *et al.* 2023b) | Pathogenicity predictor (large language model trained on human/primate variant frequency databases) | Saturation mutagenesis of SAVs (all positions) | (Cheng *et al.* 2023a) |
| CADD | dbNSFPv4 | Pathogenicity meta-predictor (sequence conservation, various other predictors) | Selected nsSNVs | (Rentzsch *et al.* 2019) |
| Condel | FannsDB | Pathogenicity meta-predictor (FATHMM, MutationAssessor) | All possible nsSNVs | (González-Pérez and López-Bigas 2011) |
| DUET | Calculation with supplied structures (web server) | Protein stability meta-predictor (mCSM, SDM) | All possible nsSNVs | (Pires, Ascher and Blundell 2014a) |
| FATHMM | dbNSFPv4 | Pathogenicity predictor (sequence homology, disease weighting) | Selected nsSNVs | (Shihab *et al.* 2013) |
| M-CAP | dbNSFPv4 | Pathogenicity predictor (sequence homology, genetic constraints) | Selected nsSNVs | (Jagadeesh *et al.* 2016) |
| mCSM | Calculation with supplied structures (web server) | Protein stability predictor (graph-based structural signatures) | Saturation mutagenesis of SAVs (all domains) | (Pires, Ascher and Blundell 2014b) |
| MetaSVM | dbNSFPv4 | Pathogenicity meta-predictor (11 other predictors, allele frequency) | Selected nsSNVs | (Dong *et al.* 2015) |
| MetaLR | dbNSFPv4 | Pathogenicity meta-predictor (11 other predictors, allele frequency) | Selected nsSNVs | (Dong *et al.* 2015) |
| MPC | dbNSFPv4 | Pathogenicity predictor (sequence homology, genetic constraints) | Selected nsSNVs | (Samocha *et al.* 2017) |
| MutationAssessor | dbNSFPv4 | Pathogenicity predictor (sequence homology) | Selected nsSNVs | (Reva, Antipin and Sander 2011) |
| MutationTaster | dbNSFPv4 | Pathogenicity predictor (sequence homology) | Selected nsSNVs | (Schwarz *et al.* 2014) |
| MutPred | dbNSFPv4 | Pathogenicity predictor (sequence homology, conservation, structural change) | Selected nsSNVs | (Pejaver *et al.* 2020) |
| MVP | dbNSFPv4 | Pathogenicity predictor (sequence conservation, structural features, other predictors) | Selected nsSNVs | (Qi *et al.* 2021) |
| PolyPhen-2 | Retrieval from web server | Pathogenicity predictor (sequence homology, structural annotation) | Saturation mutagenesis of SAVs (N2AB domains) | (Adzhubei, Jordan and Sunyaev 2013) |
| PrimateAI | dbNSFPv4 | Pathogenicity predictor (direct learning from sequence alignments) | Selected nsSNVs | (Sundaram *et al.* 2018) |
| PROVEAN | dbNSFPv4 | Pathogenicity predictor (sequence homology) | Selected nsSNVs | (Choi *et al.* 2012) |
| REVEL | REVEL genome segment files | Pathogenicity meta-predictor (13 other predictors) | All possible nsSNVs | (Ioannidis *et al.* 2016) |
| Rhapsody | Calculation with supplied structures (local pipeline) | Pathogenicity predictor (sequence conservation, structural features, coarse-grained dynamics) | Saturation mutagenesis of SAVs (N2AB domains) | (Ponzoni *et al.* 2020) |
| SIFT | dbNSFPv4 | Pathogenicity predictor (sequence conservation) | Selected nsSNVs | (Ng and Henikoff 2003) |

##### Supplementary Table 3

*Experimental structures from PDBe available in TITINdb2. The domain(s) given in the PDBe record, the corresponding UniProt domain, and the corresponding domain in the inferred complete isoform of titin given by TITINdb2 are given.*

| PDB | Reported Domain | UniProt Domain | TITINdb2 Domain | Method | Resolution |
| --- | --- | --- | --- | --- | --- |
| 2F8V | Titin-Nterm | Ig-like 1 | Ig-1 | X-Ray Diffraction | 2.75A |
|  |  | Ig-like 2 | Ig-2 |  |  |
| 2A38 | Titin-Nterm | Ig-like 1 | Ig-1 | X-Ray Diffraction | 2.00A |
|  |  | Ig-like 2 | Ig-2 |  |  |
| 6FWX | Titin-Z1 | Ig-like 1 | Ig-1 | X-Ray Diffraction | 3.00A |
|  | Titin-Z2 | Ig-like 2 | Ig-2 |  |  |
| 6SDB | Titin-Z1 | Ig-like 1 | Ig-1 | X-Ray Diffraction | 2.80A |
|  | Titin-Z2 | Ig-like 2 | Ig-2 |  |  |
| 1YA5 | Titin-Z1 | Ig-like 1 | Ig-1 | X-Ray Diffraction | 2.44A |
|  | Titin-Z2 | Ig-like 2 | Ig-2 |  |  |
| 6DL4 | Titin-Z10 | Ig-like 9 | Ig-9 | Solution NMR |  |
| 1G1C | Titin-I1 | Ig-like 10 | Ig-10 | X-Ray Diffraction | 2.10A |
| 4QEG | Titin-I10 | Ig-like 16 | Ig-19 | X-Ray Diffraction | 2.00A |
| 5JDJ | Titin-I10 | Ig-like 16 | Ig-19 | X-Ray Diffraction | 1.74A |
| 5JDD | Titin-I9 | N/A | Ig-18 | X-Ray Diffraction | 1.53A |
|  | Titin-I10 | Ig-like 16 | Ig-19 |  |  |
|  | Titin-I11 | Ig-like 17 | Ig-20 |  |  |
| 5JDE | Titin-I9 | N/A | Ig-18 | X-Ray Diffraction | 1.90A |
|  | Titin-I10 | Ig-like 16 | Ig-19 |  |  |
|  | Titin-I11 | Ig-like 17 | Ig-20 |  |  |
| 2RIK | Titin I67 | Ig-like 64 | Ig-70 | X-Ray Diffraction | 1.60A |
|  | Titin I68 | Ig-like 65 | Ig-71 |  |  |
|  | Titin I69 | Ig-like 66 | Ig-72 |  |  |
| 3B43 | Titin I65 | Ig-like 62 | Ig-68 | X-Ray Diffraction | 3.30A |
|  | Titin I66 | Ig-like 63 | Ig-69 |  |  |
|  | Titin I67 | Ig-like 64 | Ig-70 |  |  |
|  | Titin I68 | Ig-like 65 | Ig-71 |  |  |
|  | Titin I69 | Ig-like 66 | Ig-72 |  |  |
|  | Titin I70 | Ig-like 67 | Ig-73 |  |  |
| 5JOE | Titin I81 | N/A | Ig-84 | X-Ray Diffraction | 2.00A |
| 7AHS | Titin-N2A Ig81 | Ig-like 77 | Ig-84 | X-Ray Diffraction | 2.04A |
|  | Titin-N2A Ig82 | Ig-like 78 | Ig-85 |  |  |
|  | Titin-N2A Ig83 | Ig-like 79 | Ig-86 |  |  |
| 6YJ0 | Titin I83 | Ig-like 79 | Ig-86 | Solution NMR |  |
| 1TIT | Titin-I27 | N/A | Ig-94 | Solution NMR |  |
| 1TIU | Titin-I27 | N/A | Ig-94 | Solution NMR |  |
| 1WAA | Titin-I27 | N/A | Ig-94 | X-Ray Diffraction | 1.80A |
| 4O00 | Titin-A3 | FnIII 3 | Fn3-3 | X-Ray Diffraction | 1.85A |
| 8BXR | Titin I109  Titin I110  Titin I111 | N/A  FnIII 4  FnIII 5 | Ig-109 Fn3-4  Fn3-5 | X-Ray Diffraction | 2.70A |
| 8BW6 | Titin I110 | FnIII 4 | Fn3-4 | X-Ray Diffraction | 1.95A |
| 8BVO | Titin I110  Titin I111 | FnIII 4  FnIII 5 | Fn3-4 Fn3-5 | X-Ray Diffraction | 2.55A |
| 1BPV | Titin-A71 | FnIII 62 | Fn3-62 | Solution NMR |  |
| 3LPW | Titin-A77 | FnIII 66 | Fn3-66 | X-Ray Diffraction | 1.65A |
| 3LPW  3LCY | Titin-A78 | FnIII 67 | Fn3-67 | X-Ray Diffraction  X-Ray Diffraction | 1.65A  2.50A |
| 3LPW  3LCY  3LCY  2J8H | Titin-A164 | Ig-like 140 | Ig-156 | X-Ray Diffraction  X-Ray Diffraction  X-Ray Diffraction  X-Ray Diffraction | 1.65A  2.50A  2.50A  1.99A |
|  | Titin-A165 | N/A | Ig-157 |  |  |
| 3LCY  2J8H  2J8H  2J8O | Titin-A168 | Ig-like 141 | Ig-158 | X-Ray Diffraction  X-Ray Diffraction  X-Ray Diffraction  X-Ray Diffraction | 2.50A  1.99A  1.99A  2.49A |
|  | Titin-A169 | Ig-like 142 | Ig-159 |  |  |
| 2J8H  2J8O  2J8O  2ILL | Titin-A168 | Ig-like 141 | Ig-158 | X-Ray Diffraction  X-Ray Diffraction  X-Ray Diffraction  X-Ray Diffraction | 1.99A  2.49A  2.49A  2.20A |
|  | Titin-A169 | Ig-like 142 | Ig-159 |  |  |
| 2J8O  2ILL  2ILL  2NZI | Titin-A168 | Ig-like 141 | Ig-158 | X-Ray Diffraction  X-Ray Diffraction  X-Ray Diffraction  X-Ray Diffraction | 2.49A  2.20A  2.20A  2.90A |
|  | Titin-A169 | Ig-like 142 | Ig-159 |  |  |
| 2ILL  2NZI  2NZI  4JNW | Titin-A168 | Ig-like 141 | Ig-158 | X-Ray Diffraction  X-Ray Diffraction  X-Ray Diffraction  X-Ray Diffraction | 2.20A  2.90A  2.90A  2.06A |
|  | Titin-A169 | Ig-like 142 | Ig-159 |  |  |
| 2NZI  4JNW  1TKI | Titin-A170 | FnIII 132 | Fn3-132 | X-Ray Diffraction  X-Ray Diffraction  X-Ray Diffraction | 2.90A  2.06A  2.00A |
|  | Titin Kinase | Protein kinase | Kinase-1 |  |  |
|  | Titin Kinase | Protein kinase | Kinase-1 |  |  |
| 6YGN | Titin-A168 | FnIII 132 | Fn3-132 | X-Ray Diffraction | 2.40A |
| 6YGN  2BK8 | Titin Kinase | Protein kinase | Kinase-1 | X-Ray Diffraction  X-Ray Diffraction | 2.40A  1.69A |
| 6YGN  2BK8  6HCI | Titin-M1 | Ig-like 143 | Ig-160 | X-Ray Diffraction  X-Ray Diffraction  X-Ray Diffraction | 2.40A  1.69A  2.12A |
|  | Titin-M1 | Ig-like 143 | Ig-160 |  |  |
|  | Titin-M3 | Ig-like 145 | Ig-162 |  |  |
| 6H4L | Titin-M4 | Ig-like 146 | Ig-163 | X-Ray Diffraction | 1.60A |
| 3QP3 | Titin-M4 | Ig-like 146 | Ig-163 | X-Ray Diffraction | 2.00A |
| 1NCT | Titin-M5 | Ig-like 147 | Ig-164 | Solution NMR |  |
| 1NCU | Titin-M5 | Ig-like 147 | Ig-164 | Solution NMR |  |
| 1TNN | Titin-??? | Ig-like 147 | Ig-164 | Solution NMR |  |
| 1TNM | Titin-??? | Ig-like 147 | Ig-164 | Solution NMR |  |
| 3PUC | Titin-M7 | Ig-like 149 | Ig-166 | X-Ray Diffraction | 0.96A |
| 4C4K | Titin-M10 | Ig-like 152 | Ig-169 | X-Ray Diffraction | 1.95A |
| 4UOW | Titin-M10 | Ig-like 152 | Ig-169 | X-Ray Diffraction | 3.30A |
| 3Q5O | Titin-M10 | Ig-like 152 | Ig-169 | X-Ray Diffraction | 2.05A |
| 2Y9R | Titin-M10 | Ig-like 152 | Ig-169 | X-Ray Diffraction | 1.90A |
| 2WP3 | Titin-M10 | Ig-like 152 | Ig-169 | X-Ray Diffraction | 1.48A |
| 2WWK | Titin-M10 | Ig-like 152 | Ig-169 | X-Ray Diffraction | 1.70A |
| 2WWM | Titin-M10 | Ig-like 152 | Ig-169 | X-Ray Diffraction | 2.30A |
| 3KNB | Titin-Cterm | Ig-like 152 | Ig-169 | X-Ray Diffraction | 1.40A |

##### Supplementary Table 4

*Minor allele frequencies of variants reported as pathogenic in TITINdb2 from gnomAD-4.0.0, both from the global population and from reported sub-populations. A label of "." denotes variants not found in any individuals in the respective ethnic groups. Sub-population ethnicities are given as follows: AFR African; AMR Latin American; ASJ Ashkenazi Jewish; EAS East Asian; FIN Finnish; NFE European (excluding Finnish); SAS South Asian; MDE Middle Eastern. MAFs of greater than 0.0001, or 1 in every 10000 individuals, are highlighted in light green; MAFs of greater than 0.001, or 1 in every 1000 individuals, are highlighted in dark green.*

| ***Variant*** | **Global** | **AFR** | **AMR** | **ASJ** | **EAS** | **FIN** | **NFE** | **SAS** | **MDE** | **Other** |
| --- | --- | --- | --- | --- | --- | --- | --- | --- | --- | --- |
| *V54M* | 3.72E-05 | 1.33E-05 | 3.33E-05 | . | **1.11E-03** | . | 3.39E-06 | . | 1.65E-04 | 3.20E-05 |
| *A82D* | . | . | . | . | . | . | . | . | . | . |
| *A178D* | . | . | . | . | . | . | . | . | . | . |
| *E238Q* | . | . | . | . | . | . | . | . | . | . |
| *Q466R* | 6.20E-06 | 1.33E-04 | . | . | . | . | . | . | . | . |
| *R740L* | 6.84E-07 | . | . | . | . | . | 8.99E-07 | . | . | . |
| *A743V* | 1.54E-05 | . | . | . | 2.91E-04 | . | . | . | . | . |
| *W976R* | . | . | . | . | . | . | . | . | . | . |
| *V1034M* | 9.56E-04 | 1.33E-04 | 2.33E-04 | . | . | **2.56E-03** | **1.07E-03** | 6.92E-04 | 1.97E-04 | 4.81E-04 |
| *T2014A* | 3.04E-05 | . | . | . | **1.07E-03** | . | . | . | . | 1.60E-05 |
| *T2896I* | 1.59E-06 | 5.65E-05 | . | . | . | . | . | . | . | . |
| *S4116Y* | 6.16E-06 | . | . | . | . | . | 8.09E-06 | . | . | . |
| *G4714D* | 2.36E-05 | . | . | . | 6.92E-04 | . | 5.09E-06 | . | . | 1.60E-05 |
| *S4780N* | 4.40E-05 | . | . | . | **1.05E-03** | . | 3.39E-06 | 1.65E-04 | . | 8.00E-05 |
| *L4854F* | 7.69E-06 | 1.69E-05 | 5.08E-05 | . | . | . | . | 1.34E-05 | . | 2.84E-05 |
| *C5054R* | 1.20E-06 | . | . | . | . | . | 1.31E-06 | . | . | . |
| *Y9275C* | 4.96E-05 | 2.67E-05 | . | . | . | 1.56E-05 | 6.44E-05 | . | . | 1.60E-05 |
| *R9744H* | 5.27E-05 | 2.67E-05 | 5.00E-05 | . | 2.23E-05 | . | 6.10E-05 | 2.20E-05 | 1.64E-04 | 6.40E-05 |
| *R9848Q* | 1.80E-05 | . | . | . | 2.01E-04 | . | 1.27E-05 | 3.30E-05 | 1.64E-04 | 1.60E-05 |
| *A9980T* | 1.98E-04 | . | 1.67E-05 | . | **5.84E-03** | . | 1.19E-05 | 2.21E-04 | 1.65E-04 | 1.28E-04 |
| *H10092Y* | **3.49E-03** | 4.27E-04 | 7.66E-04 | 8.78E-04 | 4.46E-05 | **9.03E-03** | **3.85E-03** | **2.29E-03** | **1.65E-03** | **2.90E-03** |
| *S13702P* | 6.41E-06 | . | . | . | . | . | 1.20E-05 | . | . | . |
| *A13715E* | . | . | . | . | . | . | . | . | . | . |
| *R14640C* | 2.17E-05 | . | . | . | 2.24E-05 | . | 2.71E-05 | 1.10E-05 | . | 1.60E-05 |
| *N16133K* | . | . | . | . | . | . | . | . | . | . |
| *N16429K* | . | . | . | . | . | . | . | . | . | . |
| *W16471C* | 6.85E-07 | . | . | . | . | . | 9.00E-07 | . | . | . |
| *Y16686C* | . | . | . | . | . | . | . | . | . | . |
| *C17051R* | 1.59E-06 | . | . | . | . | . | . | 1.43E-05 | . | . |
| *L18237P* | 9.12E-04 | 1.73E-04 | 4.84E-04 | . | . | 1.09E-04 | **1.17E-03** | 1.10E-05 | 1.65E-04 | 5.61E-04 |
| *A18983T* | 1.03E-04 | 1.34E-05 | 3.34E-05 | 6.76E-05 | . | . | 1.34E-04 | . | . | 4.81E-05 |
| *P19288R* | . | . | . | . | . | . | . | . | . | . |
| *I19517T* | 1.61E-05 | . | 1.67E-05 | . | . | . | 1.53E-05 | . | 9.91E-04 | 1.60E-05 |
| *A19938T* | 3.60E-05 | 1.34E-05 | . | . | 3.59E-04 | . | 2.54E-05 | 8.79E-05 | . | 4.81E-05 |
| *N19955I* | . | . | . | . | . | . | . | . | . | . |
| *A21147T* | 4.20E-04 | 2.67E-05 | 6.70E-05 | 3.05E-04 | 2.25E-05 | 3.13E-05 | 2.12E-04 | **4.01E-03** | **1.49E-03** | 5.77E-04 |
| *A21877S* | 5.71E-05 | . | . | . | . | . | 1.78E-05 | 6.60E-04 | 9.89E-04 | 8.02E-05 |
| *V22232E* | . | . | . | . | . | . | . | . | . | . |
| *G24621R* | 7.53E-06 | . | . | . | 2.53E-05 | . | . | 1.04E-04 | . | 1.66E-05 |
| *R25480P* | 2.74E-06 | . | . | . | . | . | 3.60E-06 | . | . | . |
| *G27849V* | . | . | . | . | . | . | . | . | . | . |
| *R28118H* | 1.86E-05 | 5.33E-05 | 1.00E-04 | . | . | . | 1.61E-05 | . | . | 1.60E-05 |
| *R29293C* | 6.71E-04 | 2.67E-05 | 1.67E-05 | . | **1.88E-02** | 3.12E-05 | 8.90E-05 | 2.09E-04 | 3.30E-04 | **1.76E-03** |
| *S29303G* | 7.68E-06 | . | . | . | . | . | 1.44E-05 | . | . | . |
| *E29590Q* | 1.59E-06 | . | . | . | . | . | 2.86E-06 | . | . | . |
| *L30639P* | 1.20E-06 | . | . | . | . | . | . | 6.08E-05 | . | . |
| *P30723S* | 9.54E-05 | . | . | . | . | 1.73E-04 | 1.18E-04 | . | . | 4.84E-05 |
| *K31268T* | 3.15E-04 | 1.33E-05 | . | . | . | **1.86E-03** | 3.07E-04 | . | . | 4.32E-04 |
| *W31429R* | 1.20E-06 | . | **1.02E-03** | . | . | . | . | . | . | . |
| *P31709H* | . | . | . | . | . | . | . | . | . | . |
| *P31709R* | . | . | . | . | . | . | . | . | . | . |
| *C31712R* | 3.43E-06 | . | . | . | . | 1.89E-05 | 2.71E-06 | . | . | 1.66E-05 |
| *C31712Y* | . | . | . | . | . | . | . | . | . | . |
| *C31712W* | . | . | . | . | . | . | . | . | . | . |
| *W31729C* | 6.16E-06 | . | . | . | . | . | 8.10E-06 | . | . | . |
| *W31729L* | . | . | . | . | . | . | . | . | . | . |
| *W31729R* | . | . | . | . | . | . | . | . | . | . |
| *P31732L* | 9.30E-06 | . | 1.67E-05 | . | . | . | 1.02E-05 | 1.10E-05 | . | 1.60E-05 |
| *A31784V* | . | . | . | . | . | . | . | . | . | . |
| *N31786K* | 1.20E-06 | . | . | . | . | . | 1.31E-06 | . | . | . |
| *G31791D* | . | . | . | . | . | . | . | . | . | . |
| *G31791R* | . | . | . | . | . | . | . | . | . | . |
| *G31791V* | . | . | . | . | . | . | . | . | . | . |
| *R31847P* | 6.84E-07 | . | . | . | . | . | 9.00E-07 | . | . | . |
| *P33415L* | 3.48E-06 | . | . | . | . | . | 6.17E-06 | . | . | . |
| *R33903H* | 9.26E-05 | 4.00E-05 | . | . | **1.09E-03** | 7.83E-04 | 3.64E-05 | 1.10E-05 | 1.65E-04 | 8.01E-05 |
| *W34072R* | 2.17E-05 | . | 1.67E-05 | . | 2.23E-05 | . | 2.63E-05 | 1.10E-05 | . | 1.60E-05 |
| *V35643I* | 9.85E-05 | . | . | . | 2.01E-04 | . | 1.25E-04 | . | . | 3.20E-05 |
| *M35859T* | **1.28E-03** | 1.20E-04 | 1.83E-04 | **9.02E-03** | 2.00E-04 | 4.69E-05 | 5.76E-04 | **1.00E-02** | **6.10E-03** | **2.13E-03** |
| *T35915P* | 1.20E-06 | 6.33E-05 | . | . | . | . | . | . | . | . |
| *W35930R* | 2.74E-06 | . | . | . | . | . | 3.60E-06 | . | . | . |
| *H35946P* | . | . | . | . | . | . | . | . | . | . |
| *I35947N* | 1.12E-05 | . | . | . | . | . | 1.53E-05 | . | . | . |
| *L35956P* | . | . | . | . | . | . | . | . | . | . |
